# New perspectives in assessing environmental risks for birds: a simple TKTD framework to link growth and reproduction energy budget to chemical stress

**DOI:** 10.64898/2026.03.17.712277

**Authors:** Virgile Baudrot, Miléna Kaag, Sandrine Charles

## Abstract

Assessing the risk of pesticides to birds requires models that can extrapolate laboratory data to realistic exposure scenarios. In this work, we propose a new modeling framework BIRDkiss (Bird - Impact on Reproduction via Diet, *keep it simple and suitable*). This framework integrates a simplified Dynamic Energy Budget (DEBkiss) of organisms and the toxicokinetic–toxicodynamic (TKTD) of chemical substances according to a trait-based approach, thereby reducing the number of parameters to identify and strengthening the statistical robustness of the critical endpoints. The BIRDkiss model describes how food intake and toxicant exposure affect growth and egg production in birds over time. The model is fully embedded within an R package, including routines for calibration, validation and prediction under single-compound scenarios performed via Bayesian inference using standard data from the OECD avian reproduction tests. The BIRDkiss model also allows the simulations of scenarios under both varying food availability and multi-compound exposures based on the two classical mixture-toxicity paradigms: Concentration Addition (CA) and Independent Action (IA).

The results of calibration for single compounds show good results matching with observed weights and egg counts. From these calibrations, predictions for new exposure scenarios can be readily generated. For mixtures, the IA algorithm is simpler and does not require to scale variables as in CA. Simulations indicate that high food levels do not further increase egg production (saturation), whereas substantial food reductions markedly decrease reproduction because energy is reallocated to maintenance. Exposure to chemicals combined to low food availability amplify the decline in reproductive output.

The ready-to-use mechanistic, open-source BIRDkiss tool enables predicting the impact of pesticides on avian reproduction under realistic dietary exposure profiles. The implementation of CA and IA models is a first step toward mechanistic assessment of chemical mixtures, although validation still requires empirical mixture data. The model highlights the importance of food availability and shows that chemical stress can exacerbate the negative effects of nutritional stress. Integrating such models into regulatory frameworks could improve the ecological relevance of risk assessments.

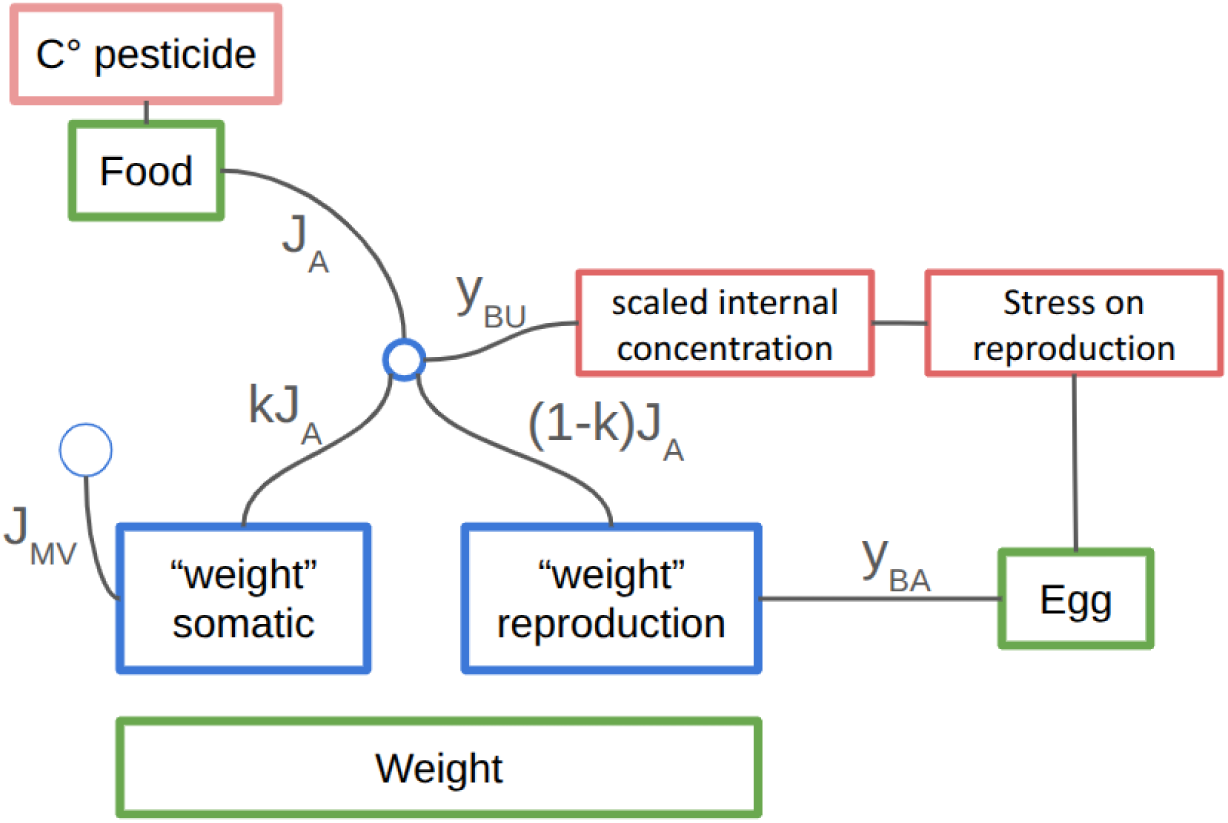

## Introduction

Environmental risk assessment (ERA) of chemicals is traditionally based on single-species laboratory testing to determine the toxicity of single active substances in formulations. However, the real challenge lies in assessing the impact of real-world scenarios involving multiple compounds and a diversity of species. Although data-driven statistical analyses are commonly employed, they are ineffective in extrapolating laboratory results to longer, time-varying exposures, such as those observed in the field. This is where mechanistic process models come in, as they can bridge this gap [Jager, 2025]. Such models are useful for predicting effects at individual and population levels, since they can be coupled with other ecosystem properties, such as environmentally realistic concentrations and population dynamics [Baudrot et al., 2021].

In this context, toxicokinetic-toxicodynamic (TKTD) models are particularly useful, as they account for the dynamic nature of toxicity and allow meaningful extrapolation from standard laboratory toxicity tests to field-relevant exposure scenarios. The European Food Safety Authority (EFSA) has recognized the added value of these models for the environmental risk assessment of pesticides [EFSA PPR Panel, 2018]. When the focus is on sub-lethal effects, such as growth and reproduction, the Dynamic Energy Budget (DEB) theory [Kooijman, 2010] allows us to consider the fitness of organisms and their life-history traits in terms of energy fluxes. This refers to the way energy, assimilated from food, is allocated to the different energy-demanding physiological functions, such as growth, maintenance, maturation, and reproduction. In a DEB-TKTD model, now recognized as fit-for-purpose in aquatic risk assessment [EFSA PPR Panel, 2018], the chemical stressors can affect the acquisition and/or the partitioning of energy. This results in different responses and/or effects on life-history traits compared to the control situation. A wide variety of TKTD models can be derived from the DEB theory, varying in complexity [Romoli et al., 2023], ranging from the simple reduced “DEBtox models” [Jager, 2024] to more complex and comprehensive DEB-TKTD models [Billoir et al., 2008]. However, only a small number of these models have been developed specifically for birds [Lamonica et al., 2025].

Avian reproduction is particularly well dedicated for such approach [Trijau et al., 2023]. First, birds are high-trophic level indicators of ecosystem health, and reproductive impairment is a sensitive, population-relevant endpoint. Second, standardized protocols such as the OECD guideline No. 206 provide high-quality data, yet current regulatory analyses often rely on descriptive statistics that fail to capture the underlying mechanistic drivers of toxicity. Developing a DEB-TKTD framework for this specific endpoint fills a critical gap in bridging standard laboratory tests with ecologically realistic risk assessment.

In addition, standard ERA frequently overlooks two key drivers of environmental risk: the presence of chemical mixtures and the inherent variability of food availability in natural settings. While single-substance testing is a regulatory cornerstone, it does not account for the interactions between chemicals [Cedergreen, 2014] or the dynamic energetic trade-offs birds face when food is limited. By integrating these complexities—mixture toxicity paradigms and food-driven energy budget dynamics—into a single framework, we aim to provide a more holistic and predictive understanding of environmental stress than current single-stressor approaches.

This paper presents a new modeling framework inspired by Trijau et al. [2023], following a simplified version of DEB models: the DEBkiss model, as proposed by [Jager et al., 2013, Jager, 2020, 2024] based on the principle of *keeping it simple and suitable*. The main difference with a standard DEB model is that there is no reserve compartment before the energy flux is split into two fractions according to the *κ* rule. In the DEBkiss approach, state variables are the masses (expressed as dry weight) of the structural body and the reproduction buffer of adults, which are easier to experimentally measure than energy fluxes. The DEBkiss has been shown to produce results similar to those of the standard DEB model when using primary parameters in both the calibration and subsequent prediction steps [Romoli et al., 2023].

Our modeling framework is applied to avian reproduction tests, as described in OECD guideline No. 206 [OECD, 1984] and is referred hereafter as **BIRDkiss**, standing for (Bird - Impact on Reproduction via Diet, keep it simple and suitable). The BIRDkiss model is fully embedded within an R-package, freely available online: https://doi.org/10.5281/zenodo.19209041. It combines a DEBkiss model with a TKTD module to assess the risk due to chemicals on the commonly measured endpoints in birds, namely the weight and the number of eggs, as a function of dietary exposure profiles that vary over time. The main advantage of this innovative modeling framework is its use of the most of the available data in the market authorization dossiers. All the algorithms for calibration, validation and prediction have been developed under a Bayesian framework.

In this study, we hypothesize that the BIRDkiss framework will reveal that chemical stress does not act in isolation but dynamically interacts with nutritional constraints. Specifically, when food availability is reduced, chemical exposure should amplify the decline in reproductive output, as organisms face a trade-off where available energy must be partitioned between maintenance and detoxification. By testing this prediction, we aim to demonstrate that the BIRDkiss model provides a more ecologically realistic risk assessment than traditional static dose-response approaches.

## Model Description

The model, fully described in Figure 1, includes a DEB component that accounts for food assimilation and energy/mass allocation, alongside a toxicokinetics process that describes how contaminated food is assimilated then converted into damage. This is followed by a toxicodynamics process, which converts the damage into a stress function applied to the DEB outputs (specifically, the energy and mass allocated to reproduction) to simulate sublethal adverse effects.

**Figure 1.**
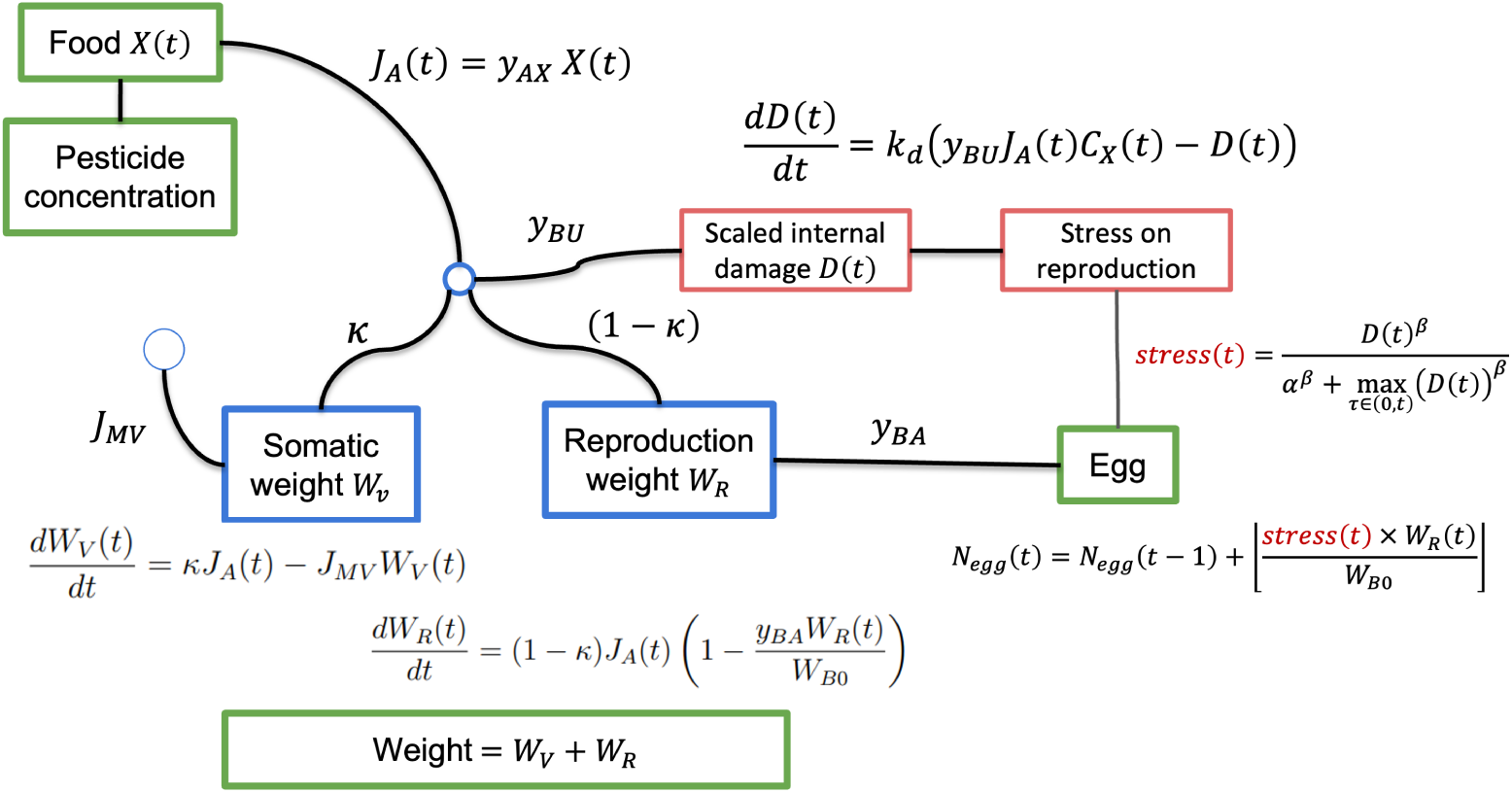
Diagram of the BIRDkiss model for the calibration, validation and prediction of avian growth and reproduction under chemical exposure via food. The green boxes represent the measured variables: the amount of food (*X_t_*), the weight of the individuals (*W_R_*(*t*) + *W_V_* (*t*)) and the production of eggs (*N_egg_*(*t*)). The assimilated food is represented by the latent variable *J_A_*(*t*), with *y_AX_* the food assimilation rate; *J_A_*(*t*) is distributed between growth and reproduction using the *κ* rule. The cost of maintenance is assumed to be linearly linked to the somatic weight through the parameter *J_MV_*. Two additional parameters influence the egg production: the egg weight, *W_B_*_0_, and the egg production cost *y_BA_*. The chemical effect is accounted for with scaled internal damage (variable *D*(*t*)) through the dominant rate constant *k_d_* (representing a compromise between accumulation and elimination) and parameter *y_BU_* (standing for bioavailability); the stress function applies to *N_egg_*(*t*) with two parameters, *α* and *β*, for a sigmoid shape.

### Dynamic Energy Budget

#### Food assimilation

The monitored amount of food provided to individuals or groups of individuals is denoted *X*(*t*) (in *g*) at each time point *t* (in days, *d*). The concentration of the chemical compound in the food items is denoted *C_X_*(*t*) (in *ppm*). Assimilation refers to the process by which ingested substances are absorbed and made available for utilization by the cells and tissues of the bird. The assimilation rate is defined by parameter *y_AX_ ∈* [0, 1] (expressed in mass of assimilates, *m_a_*, per *g* of food). The assimilated flux of food is therefore defined by *J_A_*(*t*) = *y_AX_ × X*(*t*) (expressed in *g/d*).

#### Energy allocation

The assimilated flux of food, *J_A_*(*t*), is then added to the monitored mass of the organism. To distinguish between what is allocated to reproduction and what is allocated to somatic cells and tissues (maintenance and growth), and subsequently to consider the release of egg weight, the assimilated flux of food, *J_A_*(*t*), is divided into two fractions following the *κ* rule: the growth (*κJ_A_*(*t*)) and the reproduction ((1 *− κ*)*J_A_*(*t*)) fractions.

#### Growth

The growth rate is equal to the mass allocated to soma, *κJ_A_*(*t*), minus the mass used for the structure maintenance, *J_MV_ W_V_* (*t*), as defined by the following equation:

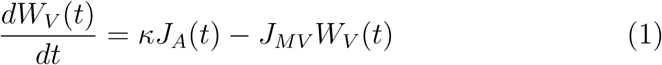

Parameter *J_MV_* represents the somatic maintenance cost, which is treated as a constant per time unit in this model, assuming a stable environment without additional stressors. However, under natural conditions, this cost should vary due to energy expenditures associated with movement (e.g., foraging, predator avoidance, migration) or internal metabolic demands (e.g., immune responses to disease or tissue repair following injury).

#### Reproduction

The reproduction mechanism underwent the most significant changes from the DEB-TKTD model proposed in Trijau et al. [2023]. The maturity maintenance cost was removed and a carrying capacity was added to the reproduction buffer to make it impossible to produce multiple eggs at the same time, without relying on a discontinuous “one egg every X hours” rule.

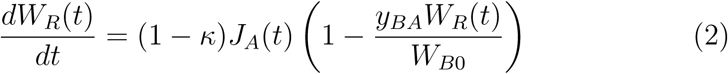

Where parameter *y_BA_* stands for the egg production cost (dimensionless), and *W_B_*_0_ for the egg weight (in grams, *g*). Then the production of eggs starts once the current time point *t* is greater than time *t_s_*, where *t_s_* is the time at which the birds start reproducing.:

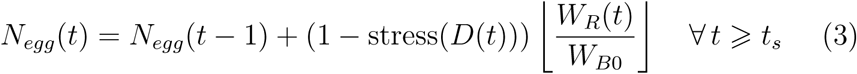

The stress function, stress(t) is neutralized for the control, where Damage (*D*(*t*) = 0). It is described later in the section dealing with chemical stress conditions. If an egg is produced, then the reproduction buffer is emptied (*W_R_*(*t*) = 0). Such a writing makes the actual cost of an egg (i.e., the mass lost from the reproduction buffer when laying occurs) vary from *W_B_*_0_ (fresh egg weight) to *W_B_*_0_*/y_BA_* (the asymptote of the reproduction buffer).

### Toxicokinetic

The proxy for the internal concentration of contaminant which may have an adverse effect but for which no measurement is available, is the scaled internal damage, denoted *D*(*t*). The damage is driven by the flux of contaminated food, which is ingested then modulated by coefficient *y_BU_* referring to bioavailability. The bioavailability is the fraction of the substance that actually enters systemic circulation (bloodstream, target tissues) and is available to exert an effect.

#### Single compound

The single-compound toxicokinetics is based on a simplified model as classically used in TKTD survival model [EFSA PPR Panel, 2018] with a single parameter *k_d_*, the dominant rate constant, which is the balanced rate between accumulation and elimination of a bioavailable chemical substance which contributes to the damage amount.

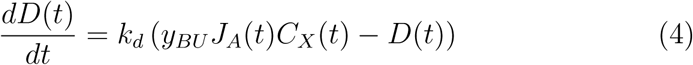

Where parameter *y_BU_*stands for the bioavailability rate.

#### Mixture of chemical substances

The only parts of the model that change in the case of multi-compound exposure are the toxicokinetics (equation (4)) and the stress function (see below). For the toxicokinetics part, equation (4) is directly reused for any compound *i* leading to write the scaled internal damage dynamics by considering the single-compound kinetic parameters *k_d,i_* and *y_BU,i_* for each compound *i* as follows:

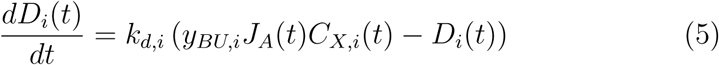

### Toxicodynamics

#### Description of the stress function

In the BIRDkiss model, the stress function, which reflects the impact of a chemical substance on reproduction, is applied to the egg production function (see equation (3)). The effect of a chemical on reducing egg production can result from many different mechanisms (e.g., neurotoxicity, disruption of hormonal pathways, or inhibition of nutrient absorption leading to nutritional deficiency). Since the observed effect is the number of produced eggs, the stress function is intended to be ”phenomenological” by modulating the egg production rate with a stress factor ranging from 0 (maximum stress; 100% reduction in egg production) to 1 (no stress; the organism produces a ”basal” number of eggs). We can also note that toxicokinetics causes the time delay between absorption and effect, along with the cumulative effect governed by excretion and metabolism rates.

#### Implementation of stress function

The stress function is consistent with the Individual Tolerance hypothesis, in which a toxicity threshold is distributed among individuals according to a log-logistic probability distribution, defined by its median, *α*, and a shape parameter, *β*, which is inversely proportional to the dispersion.

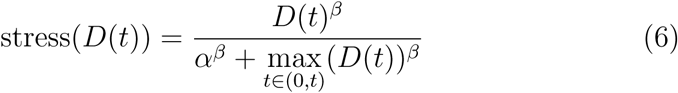

In case of multi-compound exposure, the stress function for compound *i* writes as follows:

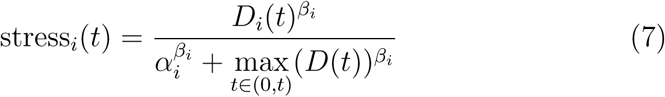

while the total resulting stress function is given by:

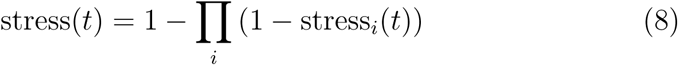

This total stress function is then used later into equation (3) driving the egg production.

### Stochasticity of model parameters and variables

Among input data, the food amount (*X*(*t*)) and the chemical concentration in food (*C_X_*(*t*)) are considered as explanatory variable and assumed to be known without error. Thus, the stochastic part of the BIRDkiss model only concerns the two following measured quantities: the total weight of an individual (or group of individuals), (*W_measured_*), and its number of produced eggs, *N_egg, measured_*.

These two variables are assumed to follow a Gamma distribution where the mean is given by the deterministic part of the BIRDkiss model (equations (1) to (2)), and the variance is controlled by a hyper-parameter:

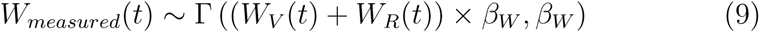

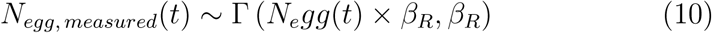

Both hyper-parameters, *β_W_* and *β_R_* were associated to non-informative priors via inverse Gamma functions with *shape* = 1 and *scale* = 0.1.

Parameters of the BIRDkiss model can be divided in two categories. One group of parameters is associated with the organism’s basal functioning and the DEB part of the BIRDkiss model (Table 1); the second group of parameters is dedicated to the effect of the specific contaminant to which organisms are exposed, i.e. the TKTD part of the BIRDkiss model (Table 2).

**Table 1.** Parameters of the DEB part of the BIRDkiss model.

| **Symbol** | **Description** | **Prior distribution** |
|---|---|---|
| *J*<sub>*MV*</sub> | somatic maintenance costs | ∼ exp(0.1) |
| *y*<sub>*AX*</sub> | assimilation rate | ∼ Beta(0.8, 0.8) |
| *y*<sub>*BA*</sub> | egg production cost | ∼ Beta(0.8, 0.8) |
| *κ* | split of assimilation flux | ∼ Beta(0.8, 0.8) |

**Table 2.**
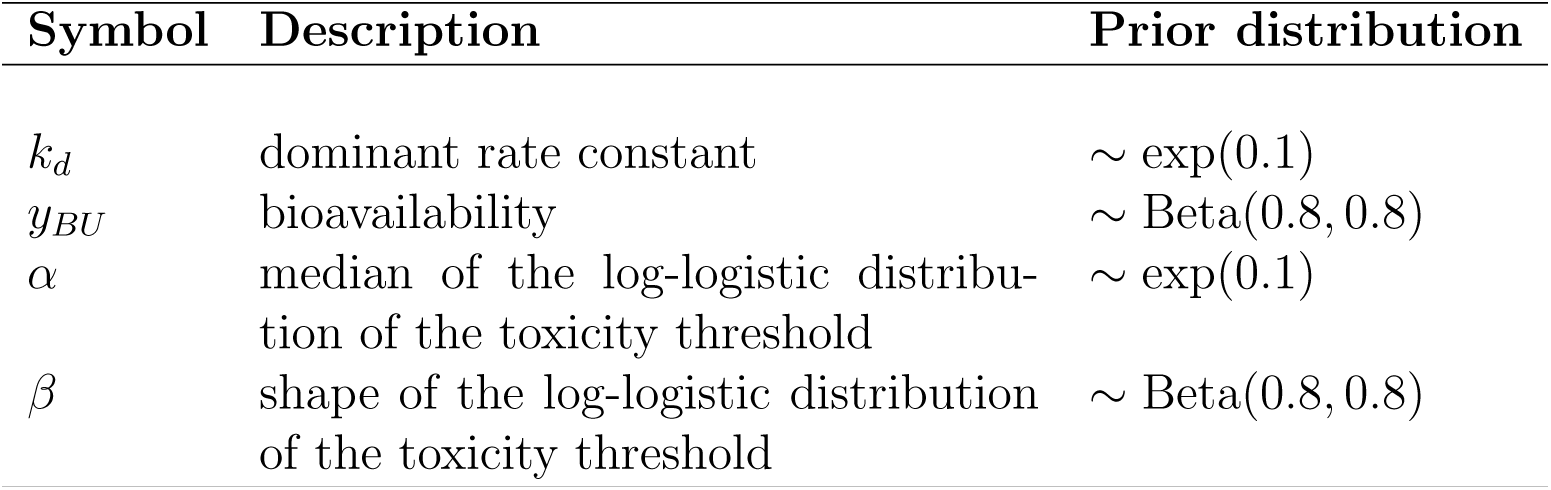
Parameters used for the TKTD part of the model.

| Symbol | Description | Prior distribution |
|---|---|---|
| *k*<sub>*d*</sub> | dominant rate constant | ∼ exp(0.1) |
| *y*<sub>*BU*</sub> | bioavailability | ∼ Beta(0.8, 0.8) |
| *α* | median of the log-logistic distribu-tion of the toxicity threshold | ∼ exp(0.1) |
| *β* | shape of the log-logistic distribution of the toxicity threshold | ∼ Beta(0.8, 0.8) |

The final BIRDkiss model only includes eight parameters, all associated to non-informative prior law distributions. All parameters ranging in [0, 1] (i.e., *y_AX_*, *y_BA_*, *κ*, *y_BU_* and *β*) have priors as Beta law distributions of shape parameters (0.8, 0.8). The cost of the somatic maintenance *J_MV_* is positive, and therefore associated to an exponential law distribution with a rate parameter equal to 0.1. We used the same parameterisation for the dominant rate constant *k_d_* of the toxicokinetics part of the BIRDkiss model. Finally, priors of parameters of the stress function, median *α* and the shape *β*, are associated with an exponential law distribution with a rate parameter of 0.1 and an inverse Gamma function with *shape* = 1 and *scale* = 0.1, respectively.

### Implementation

The BIRDkiss model is implemented in the R language [R Core Team, 2025] in combination with a Stan implementation [Stan Development, 2025] for the Bayesian inference process and the simulations. It is distributed online as an R package: https://doi.org/10.5281/zenodo. 19209041. In the R part, input data from avian reproduction tests (food intake, body weight and egg counts) are assembled into three-dimensional arrays for each replicate, producing a list that contains observation arrays, time indices and initial conditions. The Stan code defines two modules (one for inference and the other for simulation) in which six state variables (ingestion rate, body weight, reproduction buffer, damage, stress and cumulative eggs) are updated at each discrete time step using assimilation efficiencies, allocation fractions and stress parameters, and food ingestion data. Parameter are estimated under the Bayesian inference framework based on priors and the stochastic part of the model as previously defined, and predictions use the joint posterior distribution of the estimated parameters.

### Data Description

The avian reproduction test, as outlined in the OECD Test Guideline No. 206 [OECD, 1984], consists of exposing adult birds (usually mallard ducks or bobwhite quails) to a constant chemical concentration via food.

After a given period of time (previously referred as time *t_s_*), reproduction is induced through light/night exposure patterns, after which various reproduction-related endpoints are measured over time, including the number of laid eggs. The weights of the adults and their average daily food intake (approximately one measurement per week) are also recorded. In the datasets used to calibrate the model (see Table 3), the average food ingestion and egg production are typically recorded weekly, while body weights are measured once or twice during the last month of the experiment ; feeding data is averaged for each time point across replicates that share the same exposure concentration. Reproduction is induced at day 45 to 56 depending on the study and data collection continues until day 126 to 168.

**Table 3.**
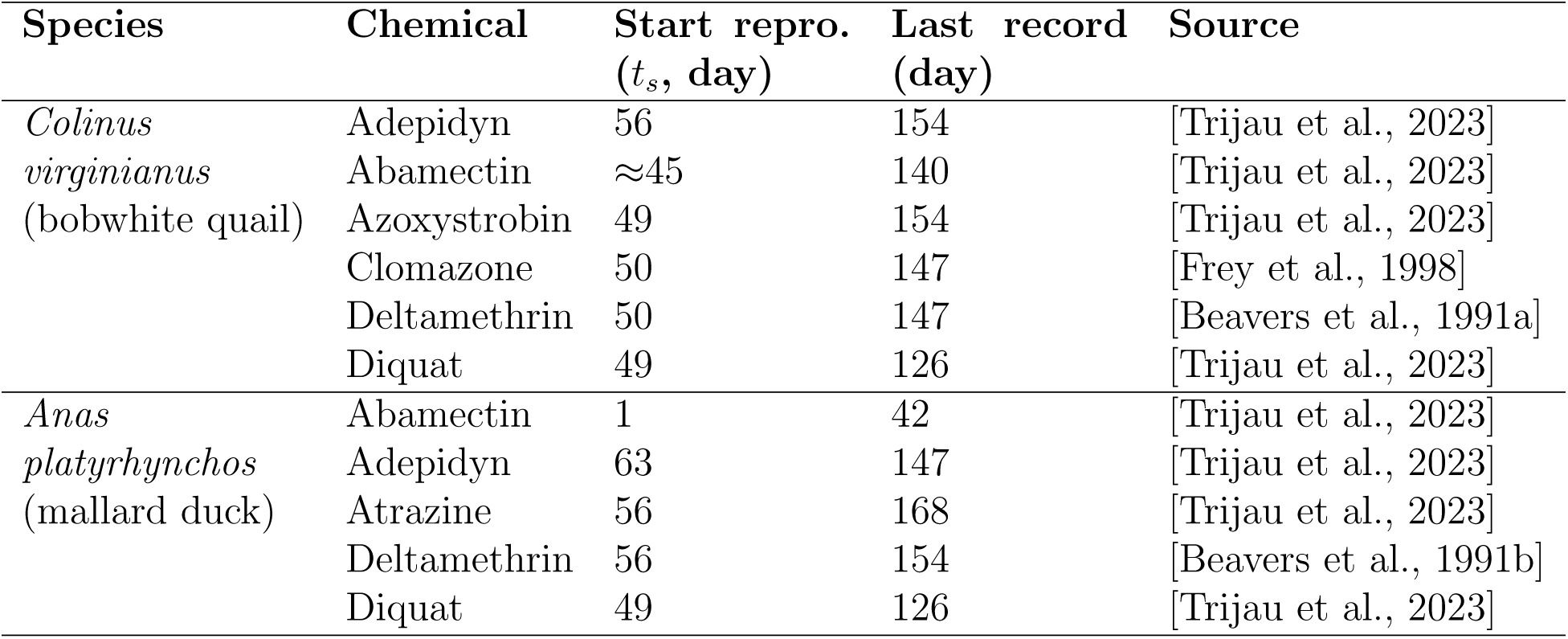
Reproduction test timelines for bobwhite quail (*Colinus virginianus*) and mallard duck (*Anas platyrhynchos*) under single chemical exposure following OECD 206 guideline.

## Results

### Calibration on single-compound

The results of calibration (Figure 2 for the duck-atrazine dataset and Supplementary Material for the other species-compound datasets) closely align with the observed data, indicating a successful calibration process that captures the biological response of ducks under various atrazine exposure conditions.

**Figure 2.**
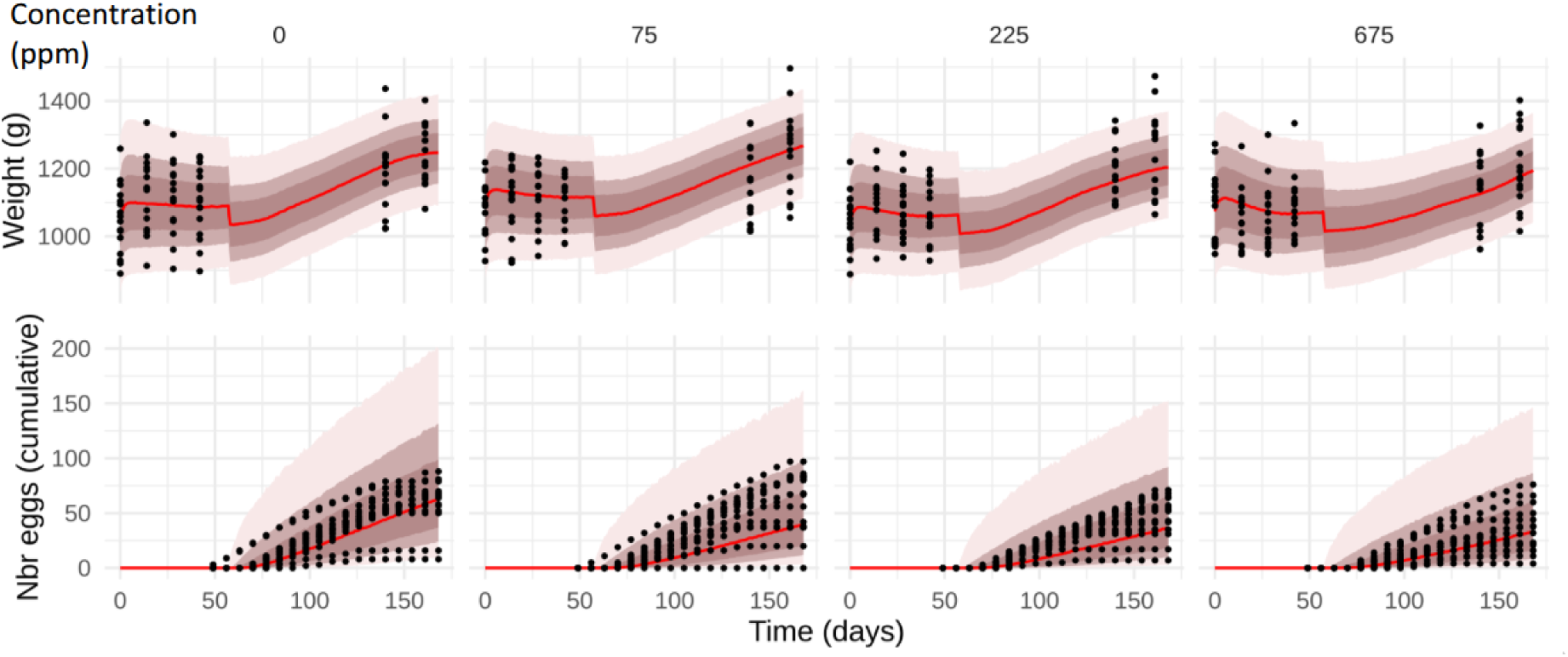
Calibration results for the mallard duck - atrazine dataset for weight (upper panel) and cumulated egg number (lower panel); each column corresponds to one exposure concentration, from 0 (left) to 675 ppm (right). The red line corresponds to the median fitted curve, while red tones stand for credible intervals at 50%, 75% and 95%.

This goodness of fit is confirmed when looking at the 95% posterior predictive check (PPC) (Figure 3 the duck-atrazine dataset, and Table 4 for the other species-compound datasets). Almost all PPC values are around 95% which confirms the effectiveness of the inference results, with model predictions adequately covering the observed data points for both body weight and cumulative egg production. Note that only two datasets (those for mallard duck exposed to abamectin and diquat) exhibit lower quality calibration results for egg production, although the prediction of weights were good in both cases. The PPC results together with the visual check of the calibration results provide strong evidence that the BIRDkiss model allows a realistic description of the biological mechanisms underlying the chemical exposure of adult bird under laboratory conditions.

**Figure 3.**
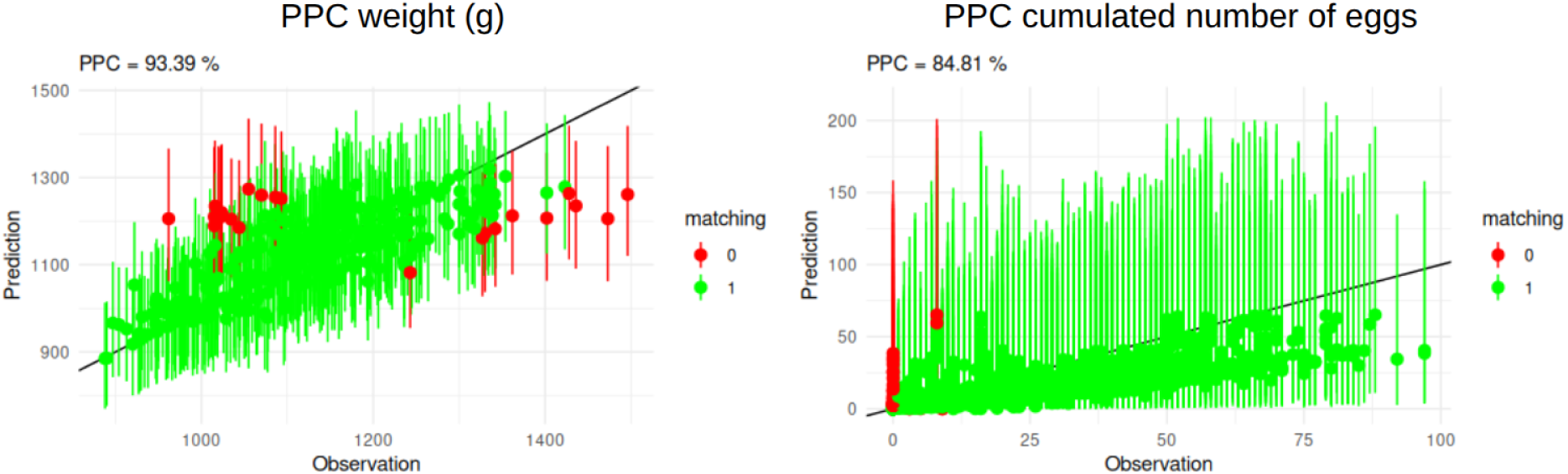
Posterior Predictive Check (PPC) computed from calibration results represented in Figure 2 for the mallard duck - atrazine dataset. PPC is computed for both weight (left panel) and cumulated egg production (right panel). The *x*-axis corresponds to observed values, the *y*-axis to the associated prediction represented as median (dots) and 95% credible interval (vertical segments). Segments appear green if the observation fall within the 95% credible interval and red otherwise. In black, the 1:1 line.

**Table 4.** Overview of calibration results for datasets on bobwhite quail (*Colinus virginianus*) and mallard duck (*Anas platyrhynchos*) under single chemical exposure with posterior predictive check (PPC) in percentage of data coverage for weights and egg productions, and the computed *EC*_50_ for egg production over 100 days of reproduction activity.

| **Species** | **Chemical** | **PPC weight (%)** | **PPC egg (%)** | *EC*<sub>50</sub> **(ppm)** |
|---|---|---|---|---|
| *Colinus virginianus*<br>(bobwhite quail) | Abamectin | 100 | 96.72 | 0.113 |
|  | Adepidyn | 98.58 | 94.44 | 114 |
|  | Azoxystrobin | 95.77 | 97.44 | 0.027 |
|  | Clomazone | 98.43 | 98.28 | 0.012 |
|  | Deltamethrin | 100 | 94.75 | 0.25 |
|  | Diquat | 95.14 | 98.96 | 1.2 |
| *Anas platyrhynchos*<br>(mallard duck) | Abamectin | 95 | **45** | 0.2 |
|  | Adepidyn | 93.75 | 100 | 250.5 |
|  | Atrazine | 93.39 | 84.81 | 1900 |
|  | Deltamethrin | 98.95 | 97.34 | 0.007 |
|  | Diquat | 92.71 | **48.96** | 18 |

In Figure 2 (see Supplementary Material for the other species-compound datasets), a sharp drop in body weight is observed around day 50, which coincides with the onset of reproduction (*t_s_*). From a biological perspective, this simulated weight loss aligns with known physiological dynamics in birds. At the onset of egg laying, the sudden and extreme metabolic demand for egg formation typically precedes the bird’s ability to sufficiently adjust its daily food intake, creating a physiological ’lag-phase’ [Kwakkel et al., 1993]. Mechanistically, the BIRDkiss model successfully captures this phenomenon through the periodic emptying of the reproduction buffer (*W_R_*) upon egg release. However, because standard OECD 206 testing protocols typically measure adult body weight only every two weeks during this critical phase, the empirical data inherently lack the fine-scale temporal resolution required to empirically verify the exact sharpness and amplitude of this biological weight drop.

### Predictions for exposure to single compounds and mixtures

The great advantage of mechanistic models such as DEB-TKTD is their ability to perform simulations for exposure profiles that vary according to (i) the amount of ingested food (e.g., food restriction conditions); (ii) variations in food contamination; (iii) both food ingestion and food contamination changes, simultaneously. This section illustrates diverse usages of the BIRDkiss model under changing environmental conditions: (1) computation of critical effect concentrations for egg production, namely *EC*_50_ values; (2) variations in both the amount of available food and the exposure concentration; (3) effects of chemical mixture; (4) effects of food restriction; and (5) joint effects of chemical mixtures and food restriction.

#### Computation of *EC*_50_ values

Table 4 shows the *EC*_50_ values for all datasets. Beyond the intrinsic interest of such concentrations leading to 50% of reduction in egg production after 100 days of exposure, these values are necessary to calculate the mixture concentration profiles under the concentration addition hypothesis (see Figure 7 for examples of scaled exposure profiles).

These *EC*_50_ values were computed by simulating the egg production with a reproduction start at day 50 and a record of the cumulative egg production at day 150, meaning 100 days of effective egg production. These simulations were performed assuming a standard food availability and exposure to a constant chemical concentration in the food items. As shown in Table 4, both species are similarly sensitive (same order of magnitude of the *EC*_50_ values) to abamectin and adepidyn. The bobwhite quail is less sensitive to deltamethrin than the mallard duck, while it is the contrary for diquat. Additionally, the bobwhite quail appears highly sensitive to azoxystrobin and clomazone, while the mallard duck is not very sensitive to atrazine.

#### Variations in food availability and chemical concentration

Although the chemical concentration in ingested food remained constant in the datasets used for calibration, the mechanistic nature of the BIRD-kiss model allows us to explore the effects of time-variable concentration profiles in food items. Figure 4 (top panel) shows four different exposure profiles via diet of adult birds and the corresponding simulations for weight (middle panel) and cumulated egg production (bottom panel). These simulations are based on the calibration results shown in Figure 2 for the mallard duck - atrazine combination.

**Figure 4.**
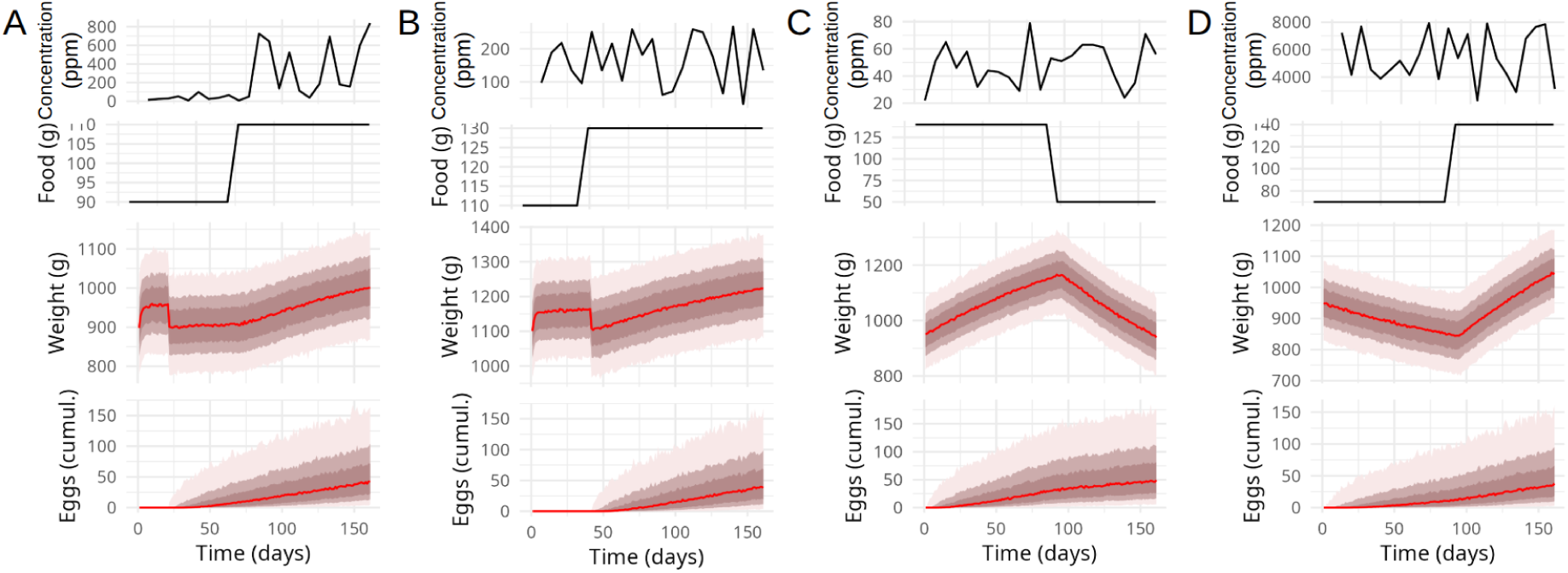
Example of predictions using the calibration results of the BIRDkiss model for the mallard duck and atrazine dataset for both weight and eggs production under different chemical exposure profiles in diet (upper panel), combined with different amounts of available food (second panel from top) and different initial conditions in terms of weight of individuals. Red lines stand for the median simulated curve, while red tones represent credible intervals at 50%, 75% and 95%.

Figure 4 illustrates the concrete usage of the BIRDkiss model for exploring realistic exposure scenarios. For instance, comparing panels A and B (Note that the *y*-axes have different scales) shows how the weight loss caused by the onset of egg production can be offset by providing additional food. Panels C and D (also with different *y* scales) illustrate the impact of food depletion on weight and, to a lesser extent, the slowdown in egg production.

#### Effects of food restriction

When an external stress is added, the BIRDkiss model can be extended by adding a new stress function to be applied on the affected latent variable. In the case of food restriction, we can fully benefit from the DEB part of the model, which describes the process of energy allocation to growth, maintenance, and reproduction from food ingestion and assimilation. A similar approach was used in [Martin et al., 2022] on rodents, with a DEB-TKTD model used to assess the effect of food variation. In particular, this study showed that it was possible to decouple a chemical direct toxic impact from its effects on the feeding rate. Because validation data was not available within our datasets, we only illustrate here the feasibility of using the BIRDkiss model to implement food restriction stress conditions and to predict their effect on weight and egg production of adult birds.

Figure 5 illustrates the sensitivity of model outcomes to significant variations in food, ranging from a 90% reduction to a theoretical 10-fold increase. More precisely, Figure 5 shows how egg production varies compared to the control group (Delta food = 1 on the *x*-axis), exhibiting a revealing trend: an excessive increase in food (right side of the *x*-axis) does not systematically lead to an increase in egg production, showing a saturation effect in the laying capacity of adult birds when food is in excess. Similarly, a substantial reduction in food availability (left side the *x*-axis) results in an even more substantial decrease in egg production, suggesting that other vital mechanisms are prioritized in resource allocation under food restriction conditions. Figure 5 even suggests an optimal zone for egg production with a 50% reduction in food availability (Delta food equal to 0.5). Of course, other considerations regarding animal welfare must be taken into account before drawing any general conclusion from these simulations.

**Figure 5.**
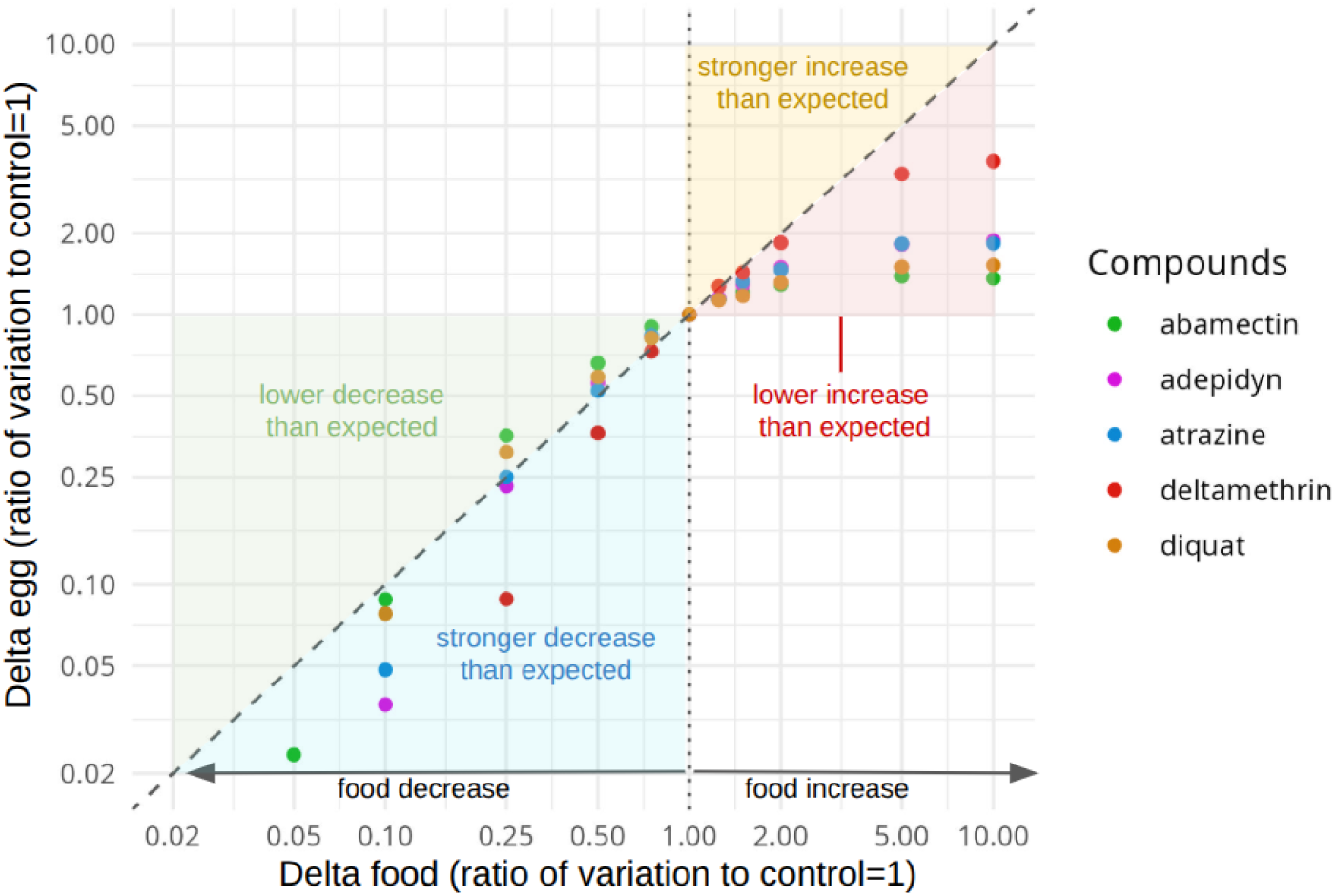
Variation in egg production as a result of variation in food availability, under control conditions of the various datasets used in this paper. The *x*-axis, Delta food, is the ratio between the food amount really ingested by adult bird and the food amount they ingest in the control data: Delta food = 1 means no variation compared to the control; 0.25 means a food amount equal to 25% of the control food amount; 2.0 means a food amount equal to 2 folds the control one. The *y*-axis is the corresponding ratio in egg production between the egg production under a changed food amount and the one of the control in the datasets. The dashed black line corresponds to a 1:1 variation, i.e., an equal % of decrease/increase in the food and in the egg production. The four colored triangles facilitate the interpretation of the dots: any dot outside these four colored triangles is unlikely as a decrease (resp. increase) in food shouldn’t imply an increase (resp. decrease) in egg production.

#### Combined effects of chemical exposure and food restriction

Figure 6 shows the variation in egg production as a function of both food amount variation and increased exposure to a chemical substance (here, atrazine; other compounds are presented in the supplementary material). The results clearly illustrate that the chemical substance amplifies the effects of reduction in food intake on the egg production. Additionally, an increase in the chemical exposure concentration does not translate into a marked effect on the egg production: on the right of the vertical blue dashed line at Delta food = 1 in Figure 6, all curves of the variation in egg production are superimposed. On the contrary, when the food amount decreases, the chemical effect becomes more and more important.

**Figure 6.**
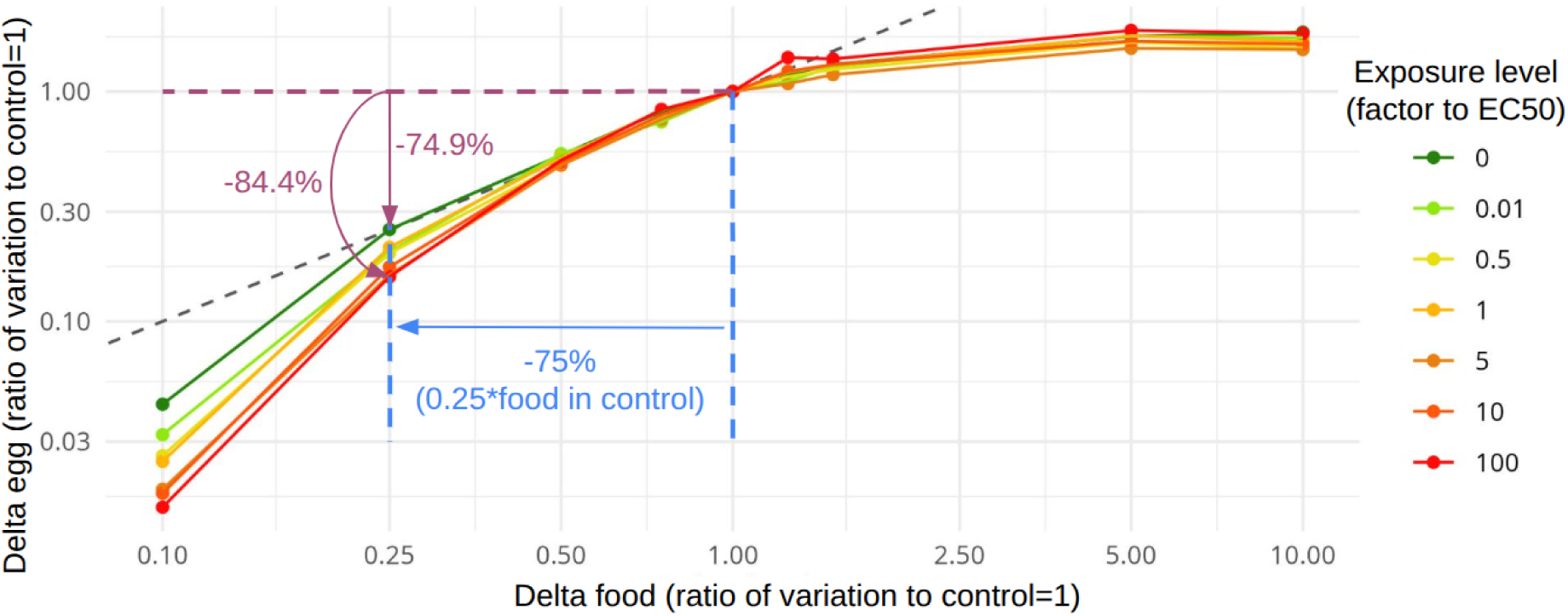
Variation in egg production as a result of a variation in both food amount and chemical exposure concentration. Results are for mallard ducks exposed to atrazine. The *x*-axis is the ratio between the given food amount and the control one. The *y*-axis is the ratio between the simulated egg production compared to the one of the control. The black dashed line corresponds to an equal variation for both food amount and egg production: a reduction of 75% in food led to a reduction of 74.9% of the egg production when the exposure concentration equals to 0, while the reduction if of 84.4% when exposure equals 100 times the *EC*_50_.

**Figure 7.**
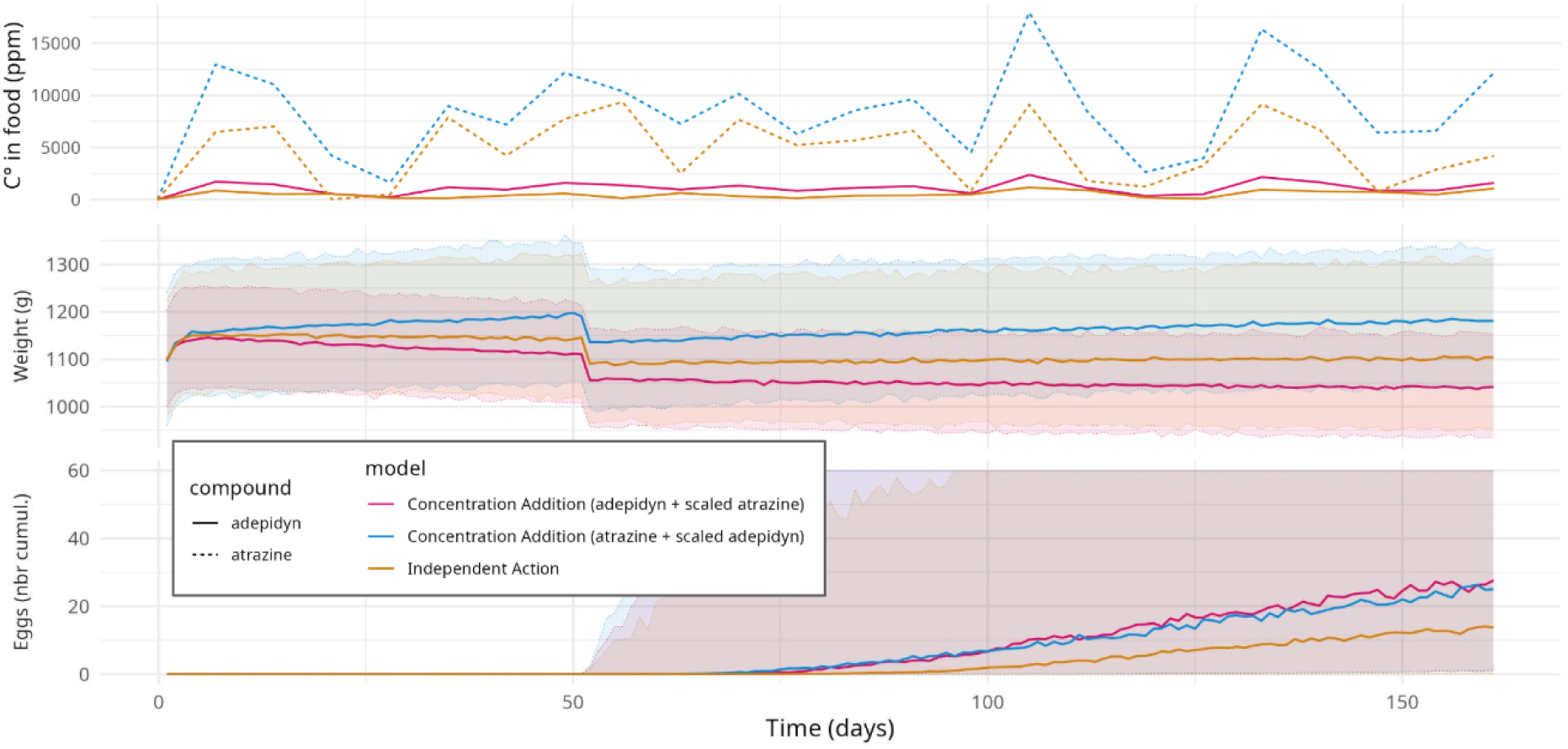
Simulation results of the BIRDkiss model for mixtures of adepidyn and atrazine based on the Concentration Addition (CA) hypothesis (pink and blue elements) and the Independent Action (IA) hypothesis (orange elements). The top panel shows the resulting exposure of a mixture of adepidyn (continuous line) and atrazine (dashed line) based on the CA hypothesis using [*B*]*_scaled_* = *^EC^*^50*,A*^ *×* [*B*] (pink and blue lines). As the IA hypothesis does not require any scaling, the model uses the “initial concentrations” (orange lines). For weight (middle panel) and egg production (bottom panel), lines represent the median predicted curve and the ribbon areas the 95% credible intervals, with colours corresponding to the exposure scenarios.

#### Exploration of chemical mixture scenarios

For chemical mixtures, the joint effects on exposed organisms are typically explained by two primary toxicity models: Concentration Addition (CA) and Independent Action (IA). Both models predict the mixture toxicity based solely on the effects of the individual compounds, assuming the absence of any toxicological interaction (synergism or antagonism) between them. Dedicated studies showed that these models correctly forecast the combined effects of pesticides in roughly 90% of cases [Belden et al., 2007, Cedergreen, 2014, Martin et al., 2021].

The CA model assumes that chemicals in a mixture behave similarly, sharing the same mode of action or influencing the same biological pathways. The total effect is anticipated by adding up the concentrations of individual chemicals, each scaled by their respective critical effective concentrations (e.g., the *EC*_50_). In practical terms, as presented in Figure 7, combining *n* chemical products under the CA model enforces to select one chemical and its calibration parameters as the reference. Each of the remaining chemicals then has its concentration scaled by an appropriate factor so that all concentrations can be pooled into the reference one. For instance, if chemical A is the reference, then the scaled concentration of B relatively to A will be 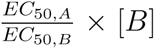, and similarly for the remaining products to be added in the mixture of interest. Then, the previous ”single-compound” toxicokinetic part of the model is used.

The IA model assumes that chemicals in a mixture act independently, with different modes of action and by affecting distinct biological pathways. The combined effect is then predicted by independently evaluating the probability of each chemical causing an effect. In practice, as illustrated in Figure 7, this implied to multiply the probabilities of egg production. For example, if one product decreases egg production by 20% and an other by 10%, their combination in a mixture will result in an egg production of 80% * 90% = 72%, that is a total of 28% reduction. This method has the advantage of avoiding the need to calculate critical effect concentrations, such as the *EC*_50_ values, to serve as a scaling factor in the application of the CA model.

The differences we found between CA and IA models (see Figure 7) are consistent with their conceptual foundations [Cedergreen, 2014]. With the CA model, the potencies of all compounds are first estimated and then summed, whereas in the IA model, the proportional responses relative to the control are multiplied. At low exposure levels, chemical substances present in the mixture below the effect thresholds for the species under study still contribute to the mixture effect in CA, while they are completely disregarded in IA. As a consequence, across the exposure scenarios presented in Figure 7, the IA model tends to yield lower combined effects than the CA model. This underestimation of the combined effects of mixture by using the IA model was systematically observed whatever the species and the chemicals mixture we investigated with the BIRDkiss model.

## Discussion

This paper presents an innovative modeling framework to link growth and reproduction energy budget of adult birds to chemical stress. Embedded within a freely available R-package, the BIRDkiss model thus offers new perspectives in assessing risk of chemical exposure combined with food limitation for birds. Combining a simplified dynamic energy budget (DEB) model to a TKTD module, the BIRDkiss model can be calibrated to standard avian toxicity test data. Also, realistic environmental scenarios with limited food amounts and exposure to multiple chemicals can be investigated in terms of effect on weight and egg production of adult birds. When calibrated on simple datasets of one bird species exposed to a single chemical substance, the BIRDkiss proved to provide relevant model parameter estimates and high quality goodness-of-fit criteria. Implemented under a Bayesian framework, the BIRDkiss allows the user to benefit of the joint posterior distribution of the parameters to predict how body weight and egg production would change under new environmental conditions, for example under time-varying chemical exposure profiles, and/or changes in food availability.

To facilitate calibration and ensure a stronger link with experimental data, the classical DEB-TKTD framework was moved towards a traitbased formulation in the BIRDkiss model, to facilitate the parametrization of the underlying processes, to ensure precise enough parameter estimates and to make the use of the model outcomes for validation relevant when comparing to standard avian toxicity test data. In addition to these methodological advances, the BIRDkiss model enables a model-guided approach to designing new or complementary experiments. As part of a de-risking strategy, the BIRDkiss model can help reduce animal testing while increasing the relevance and precision of pesticide risk assessments. Embedded within a ready-to-use tool for calibration, validation and prediction, the BIRDkiss model is a valuable asset to consider new perspectives on pesticide risk assessment for birds.

This simplified trait-based design represents a direct evolution from the previous avian DEB-TKTD framework developed by [Trijau et al., 2023]. Structurally, Trijau et al. [2023] implemented a modified standard DEB model that required an explicit reserve compartment, maturity maintenance costs, and a two-stage reproduction buffer system alongside an up-regulated assimilation flux. In contrast, BIRDkiss adopts a streamlined DEBkiss architecture [Jager et al., 2013, Jager, 2024]. This removes the explicit reserve compartment and maturity maintenance while replacing the multi-buffer system with a continuous carrying capacity on a single reproduction buffer, reducing the core parameter set to just eight primary parameters. Methodologically, while Trijau et al. [2023] relied on deterministic point-estimation (loss-function minimization evaluated with NRMSE), BIRDkiss operates within a full Bayesian inference framework using Stan. This provides joint posterior parameter distributions, explicit uncertainty quantification, and posterior predictive checks. Beyond these structural and statistical refinements for single-compound standard testing, BIRDkiss also expands the predictive scope to evaluate complex environmental scenarios by incorporating dynamic food availability (food restriction regimes) and multi-compound mixture toxicity paradigms.

Indeed, the BIRDKiss model can be used to study chemical mixtures. Both classical paradigms in mixture toxicology are implemented: the concentration addition (CA) and the independent action (IA) hypotheses [Cedergreen, 2014]. At first glance, the IA hypothesis seems more practical: it does not require the selection of a scaling factor, each compound probabilistically contributing to the global effect. Within the BIRDkiss model, the IA algorithm was simpler to implement than the CA one. Nevertheless, both assumptions lead to similar outputs, even if the IA framework tends to predict lesser effects than CA for the same conditions, as already stated in previous studies [Belden et al., 2007, Cedergreen, 2014, Martin et al., 2021]. From a regulatory standpoint, the CA hypothesis can therefore be considered the more conservative option.

The BIRDKiss simulations showed that the food availability strongly influences reproduction outputs. This is in line with the DEB models in general, which account for allocation of energy according to food assimilation. Hence, when food intake is low, the reduced allocation for repro-duction can significantly reduce the production of eggs [Jager et al., 2023]. Conversely, as shown by our simulation results, excess in food amount no longer increases egg production, meaning a saturation effect and the potential role of other limiting factors. A meta-analysis by [Ruffino et al., 2014] on avian feeding confirms that supplementary feeding increases reproduction only when natural food ingested is below a given threshold; beyond that point, the effect disappears and factors such as predation or competition limit reproduction.

The DEB-TKTD framework is particularly well suited to analyse multiple stressor patterns combining chemical exposure and food limitation. Indeed, it explicitly separate food intake and toxicity, allowing the effect of each stress factor to be distinguished and accounted for separately [Martin et al., 2022]. This separation is essential to isolate the impact of a toxicant from natural variations in food supply and to extrapolate laboratory tests to field conditions [Martin et al., 2022]. The BIRDkiss predictions highlighted a synergism between food restriction and chemical exposure. When food supply was reduced and chemical present in the exposure medium, the decline in egg production was amplified. This amplification can be explained by the fact that the available energy must be shared between maintenance and detoxification; under food limitation, this competition further reduces the energy invested in reproduction. Baas et al. [2018] noted that synergistic effects may occur when organisms face both food limitation and chemical exposure, as these stressors compete for the same energy reserves and affect both growth and reproduction. These results underline the importance of integrating abiotic and biotic factors (e.g., food, temperature) into risk assessment.

However, while BIRDkiss offers a transparent framework for linking chemical exposure to energy allocation, its applicability domain has certain limitations. First, its reliance on a simplified DEBkiss formulation omits an explicit reserve compartment [Jager, 2024], which can induce sharper numerical transitions in simulated body weight (e.g., at the onset of laying) than full DEB models [Romoli et al., 2023]. Second, the somatic maintenance costs are modeled as constant. While suitable for standard laboratory tests (e.g., OECD 206 [OECD, 1984]), this assumes a stable environment and ignores variable metabolic demands (e.g., foraging, thermoregulation) encountered by wild birds. Third, the toxicodynamic stress function remains phenomenological, modulating egg production without resolving specific physiological modes of action (e.g., endocrine disruption). Finally, calibration is currently bounded by the sparse temporal resolution of standard testing protocols (e.g., bi-weekly weighing).

Consequently, the current applicability domain is centered on adult birds under controlled testing regimes; extrapolating BIRDkiss to highly dynamic field conditions or alternative life stages requires careful consideration of these structural simplifications.

Indeed, for wild species, the food availability is likely more critical than for captive individuals used in standard avian toxicity tests. In natural settings, organisms are often exposed to harsher or more variable conditions, where the combined stress of contamination and food restriction can therefore lead to amplified effects. This is what the BIRDkiss allows to investigate. This suggests that predictions based on calibrations from solely laboratory toxicity data could underestimate the severity of impacts in free-living populations, highlighting the need to account for ecological realism when extrapolating model results.

## Conclusion

The BIRDkiss model proves to be a simple but efficient model to provide robust calibration and prediction results on the effects of single chemical substances and mixtures on bird reproduction, under food limitation or not. Made available as an open-source platform, it offers new perspectives in assessing environmental risks for birds. The flexibility of the Bayesian framework allows new data to be easily incorporated and calibration results to be generated together with estimated uncertainties. If validation for chemical mixtures still requires specific experimental data, integrating the BIRDkiss model into regulatory procedures (e.g., as the recommended data analysis in the OECD guideline 206) could lead to more realistic risk assessment.

## Supporting Information

All scripts and data are available at: https://doi.org/10.5281/zenodo. 19209041

## Acknowledgments

We thank Marie Trijau and Simon Hansul for their help in curating the data and deeply analyzing first versions of the model. This work was performed under the umbrella of the French GDR “Aquatic Ecotoxicology” framework which aims at fostering scientific discussions and collaborations for more integrative approaches in ecotoxicology.

## Credit authorship contribution statement

**Conceptualisation**: VB, MK, SC ; **Data Curation**: MK, VB, SC ; **Formal Analysis**: MK, VB, SC ; **Funding Acquisition**: SC ; **Methodology**: VB, MK, SC ; **Project Administration**: SC ; **Software**: VB, MK ; **Supervision**: SC, VB ; **Interpretation & Validation**: VB, MK, SC ; **Visualisation**: VB, MK ; **Writing – Original Draft Preparation**: VB, MK, SC ; **Writing – Review & Editing**: VB, MK, SC.

